# Sequencing accuracy and systematic errors of nanopore direct RNA sequencing

**DOI:** 10.1101/2023.03.29.534691

**Authors:** Wang Liu-Wei, Wiep van der Toorn, Patrick Bohn, Martin Hölzer, Redmond Smyth, Max von Kleist

## Abstract

Direct RNA sequencing (dRNA-seq) on the Oxford Nanopore Technologies (ONT) platforms can produce reads covering up to full-length gene transcripts while containing decipherable information about RNA base modifications and poly-A tail lengths. Although many published studies have been exploring and expanding the potential of dRNA-seq, the sequencing accuracy and error patterns remain understudied. We present the first comprehensive evaluation of accuracy and systematic errors in dRNA-seq data from diverse species, as well as synthetic RNA. Deletions significantly outnumbered mismatches/insertions, while the median read accuracy exhibited species-level variation. In addition to homopolymer errors, we observed systematic biases across nucleotides and heteropolymeric motifs in all species. In general, cytosine/uracil-rich regions were more likely to be erroneous than guanines/adenines. Moreover, the systematic errors were strongly dependent on local sequence contexts. By examining raw signal data, we identified underlying signal-level features potentially associated with the error patterns. While read quality scores approximated error rates at base and read levels, failure to detect DNA adapters may lead to data loss. By comparing distinct basecallers, we reason that some sequencing errors are attributable to signal insufficiency rather than algorithmic (base-calling) artefacts. Lastly, we discuss the implications of such error patterns for downstream applications of dRNA-seq data.

## 1 Introduction

Envisioned in the 1980s and developed over the next three decades, the nanopore sequencing technology has transformed the DNA and RNA sequencing landscape in recent years [Deamer et al., 2016, Marx, 2023]. Sequencing platforms released by the Oxford Nanopore Technologies (ONT) have enabled high-throughput long-read sequencing of single DNA/RNA molecules with low experimental requirements. Recently, it was reported that DNA reads from the latest R10.4 chemistry achieved comparable sequencing accuracy on bacterial genomes without having to be polished with short reads, thanks to significant improvements in the accuracy of homopolymer regions [Sereika et al., 2022].

In addition to DNA sequencing, ONT offers currently the only commercial platform for the direct sequencing of RNA molecules (dRNA-seq) [Garalde et al., 2018, Jain et al., 2022]. Ligated to a DNA adapter containing a helicase motor, a polyadenylated RNA molecule is translocated in the 3’–5’ direction through a protein nanopore embedded in an electrically charged membrane. The translocation causes systematic disruptions to the ionic current flow, characteristic of the nucleotides passing through the pore at the time. The current signals are basecalled into nucleotide sequences with machine learning algorithms, commonly known as “basecallers”. Ideally, these basecallers are trained on diverse organisms for better generalization and fewer biases to rare sequences and species.

Traditionally, high-throughput RNA sequencing protocols have relied on reverse transcription and/or amplification, which introduce various errors and biases that confound downstream analyses [Hansen et al., 2010, Schulz et al., 2021]. In comparison, nanopore dRNA-seq avoids these biases and at the same time, produces reads up to thousands of bases in length. The ability to cover full-length gene transcripts at single read resolution can significantly improve on analyses traditionally complex with short read RNA-seq, such as identifying transcript isoforms and quantifying Poly-A tail lengths [Workman et al., 2019, Soneson et al., 2019, Chen et al., 2021]. So far, nanopore dRNA-seq has been applied to diverse organisms such as DNA and RNA viruses [Depledge et al., 2019, Viehweger et al., 2019, Kim et al., 2020, Price et al., 2020], bacteria [Grünberger et al., 2022a], archaea [Grünberger et al., 2022b], plants [Parker et al., 2020, Rousseau-Gueutin et al., 2020, Gao et al., 2021b], yeast [Liu et al., 2019, Jenjaroenpun et al., 2021], fish [Begik et al., 2022a], mouse [Bilska et al., 2020] and humans [Workman et al., 2019, Hendra et al., 2022].

Another promising application of dRNA-seq is the characterisation of the “epitranscriptome”. Although over 100 types of RNA modifications are known, only a small subset of them, such as m^6^A, m^5^C, and pseudouridine, are transcriptome-wide detectable [Mattick and Amaral, 2023], albeit with limited accuracy [Grozhik and Jaffrey, 2018]. By sequencing RNA molecules directly, dRNA-seq has the potential to produce characteristic signals not only for the four canonical RNA bases but also for the diverse family of RNA base modifications. So far, several functionally important RNA modifications are found to be detectable with dRNA-seq, with a heavy focus on m^6^A [Liu et al., 2019, Pratanwanich et al., 2021, Gao et al., 2021a, Hendra et al., 2022, Price et al., 2020], but also pseudouridine [Begik et al., 2021] and inosine [Nguyen et al., 2022]. Typically, modifications are detected by computationally identifying systematic deviations in signal levels or basecalling errors at potentially modified positions. This is typically performed by including un-modified control samples (gene knockouts or *in vitro* transcribed RNA) as baseline, due to the abundance of background errors even in unmodified RNAs.

Despite the growing interest in this technology, nanopore dRNA-seq has been widely perceived as error-prone, with reported read accuracies of around 90% for single species [Rousseau-Gueutin et al., 2020, Jain et al., 2022, Grünberger et al., 2022a], which hinders the downstream applications (by requiring control samples or short-read polishing) and limits its wider popularisation. A systematic examination and characterisation of dRNA-seq accuracy and error patterns could help 1) clarify its current limitations and potential, 2) direct future computational methods to account for the underlying uncertainty/errors, and 3) establish the foundation for improving pore chemistry and base-calling algorithms. Here, starting from the raw signal data, we re-analysed twelve public datasets covering a wide taxonomic range and present the first comprehensive benchmark of dRNA-seq accuracy using standardized metrics. Notably, systematic errors exist at both single base and motif levels that are reproducible across all investigated organisms, and such errors show strong and complex dependency on their local sequence contexts. In addition, we examined the relationship between read quality scores and error rates, and how adaptor detection failure can impact the read quality of short sequences. Lastly, we discuss the implications of systematic sequencing errors in downstream analyses of dRNA-seq, such as on transcript isoform identification and base modification detection.

## 2 Methods

### 2.1 Data

We collected public dRNA-seq datasets covering a diverse range of organisms and *in vitro* transcribed RNAs, sequenced with the ONT direct RNA kits SQK-RNA001 and SQK-RNA002. Sequencing datasets in raw *fast5* files from the following studies are included in the analysis of sequencing accuracy and errors:

1) native and *in vitro* transcribed human data [Jenjaroenpun et al., 2021],

2) the house mouse *Mus musculus* [Bilska et al., 2020],

3) yeast wildtype strain SK1 [Liu et al., 2019],

4) *Escherichia coli* [Grünberger et al., 2022a],

5) in-cell and *in vitro* transcribed SARS-CoV-2 [Kim et al., 2020],

6) *Caenorhabditis elegans* [Roach et al., 2020],

7) the zebrafish *Danio rerio* [Begik et al., 2022a],

8) *Arabidopsis thaliana* [Parker et al., 2020],

9) *Haloferax volcanii* [Grünberger et al., 2022b],

10) *in vitro* transcribed short RNAs [Begik et al., 2021], from a) *Bacillus subtilis* guanine riboswitch, b) *B. subtilis* lysine riboswitch, and c) *Tetrahymena* ribozyme (reference length 202, 274 and 460 bases, respectively). The reference sequences for organisms 1), 2), 4), 6), 7), 8), 9) are obtained from the NCBI reference genome database and are respectively Human Build 38 GRCh38.p14, Mouse Build 39 GRCm39, *E. coli* ASM584v2, *C. elegans* WBcel235, Zebrafish Build 11 GRCz11, *A. thaliana* TAIR10.1 and *H. volcanii* DS2 ASM2568v1, while the references for 3), 5) and 10) are provided by the original authors on the respective GitHub repositories.

### 2.2 Basecalling, mapping and filtering

Each dataset consists of raw *fast5* files that were base-called with the ONT basecaller Guppy (version 6.1.7) using the RNA model `rna_r9.4.1_70bps_hac`, with the option `--disable_qscore_filtering` to switch off the default read Q-score filtering threshold of 7.

The basecalled reads in *fastq* format are aligned to the respective reference genomes with minimap2 [Li, 2018] version 2.24, with the follow-ing options `-ax splice -k14 -uf -secondary=no --eqx --sam-hit-only`, which allows for split read alignment and disables secondary alignments. Aligned reads are further filtered to exclude supplementary alignments and reads with a mapping quality score (“MapQ”) lower than 20 or shorter than 100 bases.

To compare Guppy with the community-published basecaller RODAN [Neumann et al., 2022], we used the pretrained model provided at its GitHub repository (https://github.com/biodlab/RODAN). As RODAN failed to process some of the older versions of single read *fast5* files for some datasets, we converted them into multi-read *fast5* files using the `single_to_multi_fast5` function provided in the `ont_fast5_api` library (https://github.com/nanoporetech/ont_fast5_api).

### 2.3 Evaluation metrics

After alignment and filtering, the alignment errors (mismatches, insertions and deletions) in the reads are counted by parsing through the CIGAR string in the `.sam` file format. For the accuracy of a single read, we used the following definition

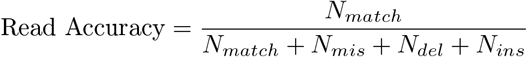

where *N*_*match*_, *N*_*mis*_, *N*_*del*_ and *N*_*ins*_ refer to the number of bases on a read that are reported as match, mismatch, deletion and insertion by the aligner. Similarly, the mismatch, insertion and deletion rate per read is defined as

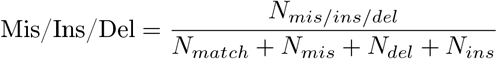

### 2.4 Systematic and context-dependent errors

To evaluate systematic errors at single nucleotide and motif level, we parsed the CIGAR string of the aligned reads from minimap2 and constructed the error profiles, which are deletions and mismatches for single nucleotides, and additionally insertions that occur within 2-mer motifs. Larger datasets are subsampled to 200,000 reads each and reads aligned to soft–masked positions (represented as lowercase letters) and ambiguous positions (represented as N/n) in the reference genomes are excluded.

For the analysis of homopolymers, the accuracy of a homopolymer motif is defined as the frequency of the homopolymer being correctly basecalled (as the correct nucleotide type of the correct length). The base-called length of a homopolymer is counted only when the basecalled sequence is a homopolymer of the correct nucleotide type.

For the analysis of context-dependent errors, the “center error rate” was computed as the frequency of the incorrect basecalls of the center nucleotide within a 3-mer context where the first and last bases are correctly sequenced, in order to limit the effect of potential long-range errors propagating through the neighboring positions.

### 2.5 Signal analysis

The signal event data in the basecalled *fast5* files outputted by Guppy were extracted using the `ont_fast5_api` library. Guppy first determines the start signal position of basecalling (named `first_sample_template`). The raw signals are first segmented with a chunk size of ten observations each and then the transitions between nucleotides are determined by Guppy and stored in the “Move” table. Therefore, the length of raw signals of a read is roughly `first_sample_template` + 10 * the length of Move. The mean signal intensities and standard deviations are computed from the signal segments assigned to a base at only the correctly basecalled positions (based on alignment). The dwell times are computed as the length of the signal segments assigned to a base (and thus are multiples of 10).

To plot the signals of a 3-mer motif, due to the differences in dwell times, a running window is applied to extract the mean signal intensity with a step size of length of signals divided by 10 (and thus resulting in ten signal samples for each position). Only the correctly basecalled 3-mers at all three positions are included.

### 2.6 Read quality score

In addition to the per-base quality scores typically found in *fastq* files, Guppy also outputs a quality score per read, “read Q-score”, which is reported as `mean_qscore_template` in the `sequencing_summary.txt` file. Q-scores are typically used as a filter for poorly sequenced and basecalled reads. Guppy, by default, classifies reads with a Q-score lower than 7 into the `failed` folder, which is usually excluded in downstream analysis.

As the quality scores are calibrated to follow the expected error distribution of Phred quality scores [Cock et al., 2010], the expected error probability of a read is then

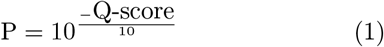

where P is essentially 1 - Expected Read Accuracy. Consequently, Guppy’s default read filter at a Q-score of 7 corresponds to an expected error rate of 20%, or equivalently, a read accuracy of 80%.

Alternatively, the read Q-score can be obtained from *fastq* files by calculating the mean per-base error rate, followed by a back conversion to Phred Q-score:

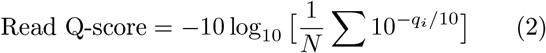

where *q*_*i*_ is the individual base quality for base *i* in a read of length *N*.

### 2.7 Adapter detection analysis

For dRNA-seq, RNA molecules are ligated to a 3’ DNA sequencing adapter. As a result, the start of the read contains signals from the DNA adapter. Guppy has built-in capabilities to detect the end of the DNA adapter and begin basecalling only at the start of the RNA molecule. The start of basecalling is reported as `first_sample_template` in the *fast5* files and used in our analysis.

## 3 Results

### 3.1 Deletions outnumber mismatches and insertions

We downloaded publicly available dRNA-seq datasets in raw *fast5* format consisting of both native and *in vitro* transcribed (IVT) samples of diverse organisms. The raw signal data were then basecalled into reads with the ONT basecaller Guppy, without the default read quality filtering. The basecalled reads were aligned to the respective references and further filtered by alignment quality and a minimum length threshold of 100. The statistics regarding each dataset included in the evaluation are given in Supplementary Table S1. The datasets range from 22,315 aligned reads in the archaeon *H. volcanii* to over 2.3 million aligned reads in the *in vitro* transcribed SARS-CoV-2 sample. For most organisms, the median read accuracy is within 88%–92%, with the mouse and the zebrafish datasets being the least accurate at 87.8% and 86.7%, respectively. The median read accuracy of 90% is in general agreement with previous reports for dRNA-seq transcriptomes of single organisms [Rousseau-Gueutin et al., 2020, Jain et al., 2022, Grünberger et al., 2022a]. The distributions of mismatches, insertions and deletions are similar across most organisms, where deletions represent the most frequent type of error at around 5% per read, while mismatches and insertions each account for 2–3% (Figure 1a). The reads from the IVT human sample have overall 1.5% fewer errors than the native human sample, possibly due to the absence of RNA base modifications. However, such a difference is not seen between the two SARS-CoV-2 samples.

**Figure 1:**
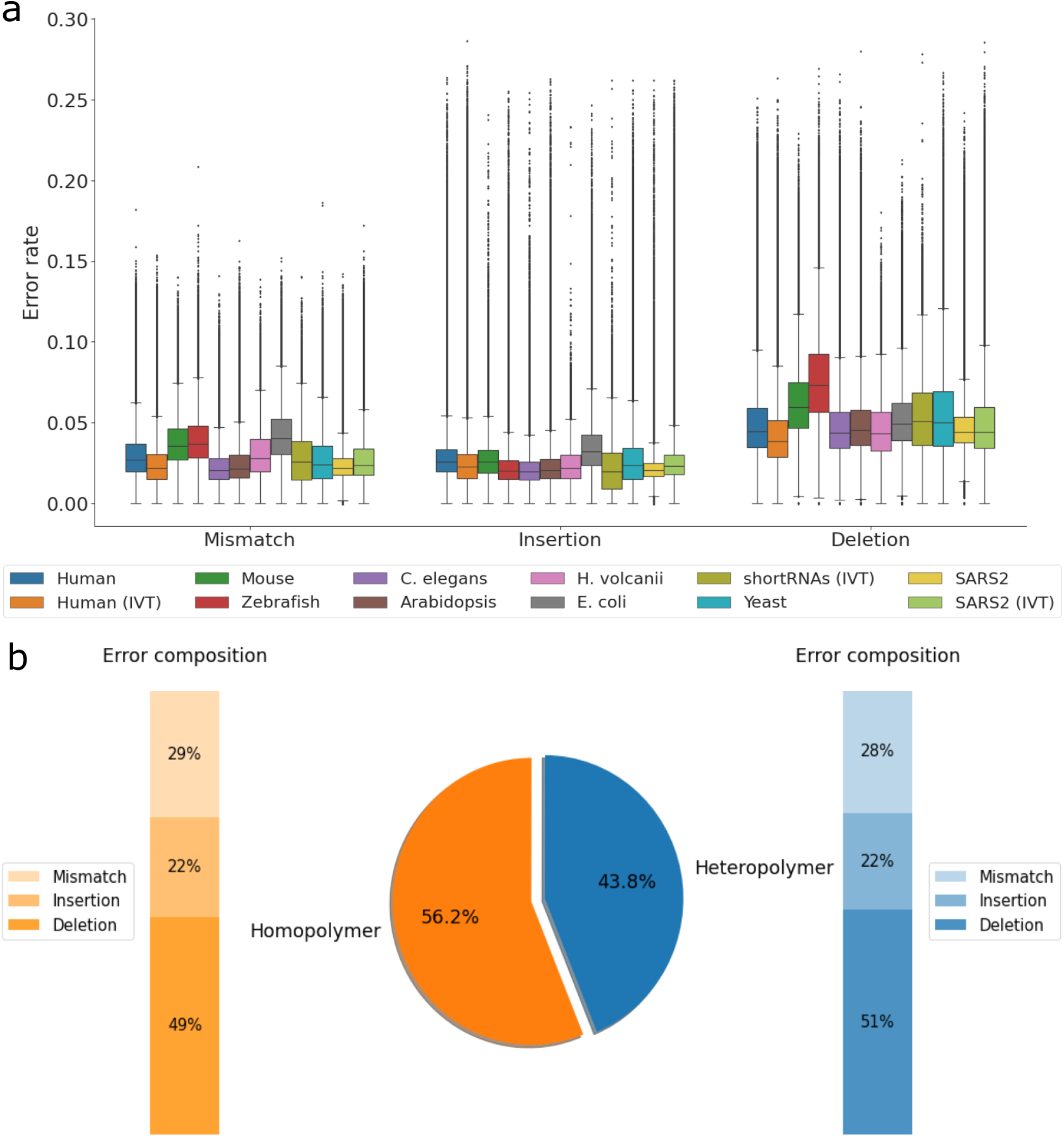
Sequencing accuracy of nanopore dRNA-seq. a) An overview of sequencing accuracy across diverse organisms, regarding mismatch, insertion and deletion rate per read. Reads with accuracy lower than 70% were filtered for visualization. IVT – *in vitro* transcribed RNA. b) The relative distribution of all errors in homopolymeric and heteropolymeric regions, based on the native human dataset. The center pie chart shows the overall proportion of errors and the respective bar plots show the composition of each error type.

While the majority of errors occur in homopolymeric regions (motifs consisting of repeats of one nucleotide), more than 40% of the errors in dRNA-seq still arise in heteropolymeric regions (Figure 1b, native human data). Moreover, the relative distribution of each error type (mismatches and indels) is similar between homopolymers and heteropolymers, with half of the errors being deletions.

To investigate whether the lengths of reads are related to their accuracy, we grouped reads by length, and found that the median read accuracy remains roughly constant as read lengths increase (Supplementary Figure S1). However, the distribution of read accuracy tends to have a larger variance in shorter reads.

### 3.2 Systematic errors in single nucleotides and heteropolymers

To explore how the sequencing errors are distributed at single nucleotide level, we computed the frequency of (in)correct basecalls across the four RNA bases (Figure 2a, Supplementary Figure S2). Across all datasets, guanines have consistently the highest accuracy, whereas cytosines and uracils tend to be the least accurate. Similar to the accuracy at read level, deletions remain the most frequent type of error at the individual base level, but are much more prevalent in Cs and Us. In terms of mismatches, C bases are more likely to be basecalled as U while G bases are more often confused with A.

**Figure 2:**
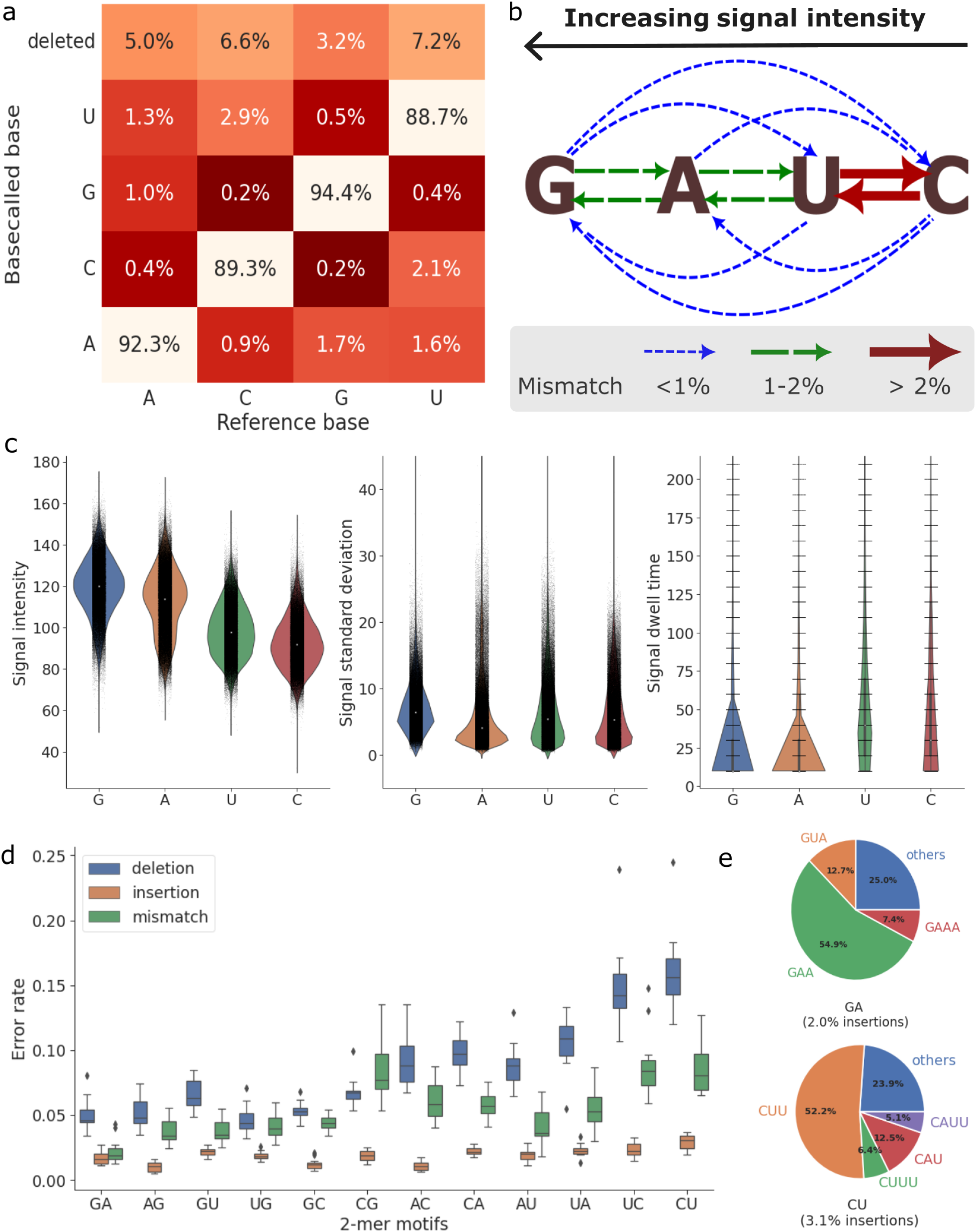
Systematic errors in nanopore dRNA-seq. a) Confusion matrix showing the frequencies of each base being correctly basecalled, miscalled or deleted, computed by taking the mean of all samples. b) The relationship between signal similarity of nucleotides and their mismatch profiles. The arrow types indicate the mismatch rate from one nucleotide to another. c) The distributions of signal features (intensity, standard deviation and dwell time) at correctly basecalled positions, grouped by nucleotide type. d) The error rate of 2-mer motifs across the samples, grouped by error type. Each data point corresponds to the accuracy of the motif in one dataset. e) The insertion error profiles of GA and CU motifs, based on the native human dataset. Inserted motifs fewer than 5% of the total number of insertions are grouped into “others”.

The mismatch error patterns are consistent with the similarity of the raw signals between the bases (Figure 2b), where the four bases ranked by signal intensity are G > A > U > C (Figure 2c left panel, thus C is most similar to U). Mismatch errors between neighboring nucleotides (especially U and C) in the signal space are generally more frequent than mismatch error rates with nucleotides that are further apart in signal space. Moreover, while the variance of signals are similar across nucleotides, the dwell times (the number of raw signal observations assigned to a single basecall by Guppy) of G and A bases are considerably shorter than U and C bases (Figure 2c). The difference in dwell times could be possibly related to the larger amount of deletion errors of U and C bases, for which the base-caller may require more information to correctly determine the number of bases.

Similarly, we found large differences between the errors of length-2 heteropolymeric motifs, especially for deletions and mismatches (Figure 2d). Among the 2-mers, GA/AG are the most accurate while UC/CU have consistently the lowest accuracy. Interestingly, although most 2-mers are close to their mirrored counterparts (“XY” and “YX”), there exist clear differences between certain pairs. In particular, GC motifs have consistently higher accuracy than CGs. Whilst deletions were generally more common than mismatches and insertions, for some 2-mers mismatch errors were very close to the deletion rate, or even higher as seen in the “CG” motif. One possible explanation for the lower deletion error in some 2-mers is that these motifs are more different in signal space, and as a consequence more distinguishable by the basecaller. Insertion errors were similar for all 2-mers, suggesting that insertions likely arise from signal noise that is not motif-specific. The inserted bases generally consist of repeats of the second base in the motif, but can also occasionally involve other bases, for example, GA → GUA (Figure 2e, Supplementary Figure S3).

### 3.3 Systematic errors in homopolymers

Accurately sequencing and basecalling homopolymers (motifs consisting of repeats of one nucleotide) is a well-known challenge in nanopore DNA sequencing [Delahaye and Nicolas, 2021, Sereika et al., 2022]. Unsurprisingly, we observed similar declines in the accuracy of homopolymer motifs in the dRNA-seq datasets as the motif length increases, especially in homopolymers longer than four bases (Figure 3a). In addition to the decreasing accuracy, the basecalled lengths of homopolymers are also more likely to be underestimated with increasing lengths (Figure 3b). While the lengths of most homopolymers of length 2 and 3 are basecalled correctly, homopolymers longer than 4 bases are consistently underestimated.

**Figure 3:**
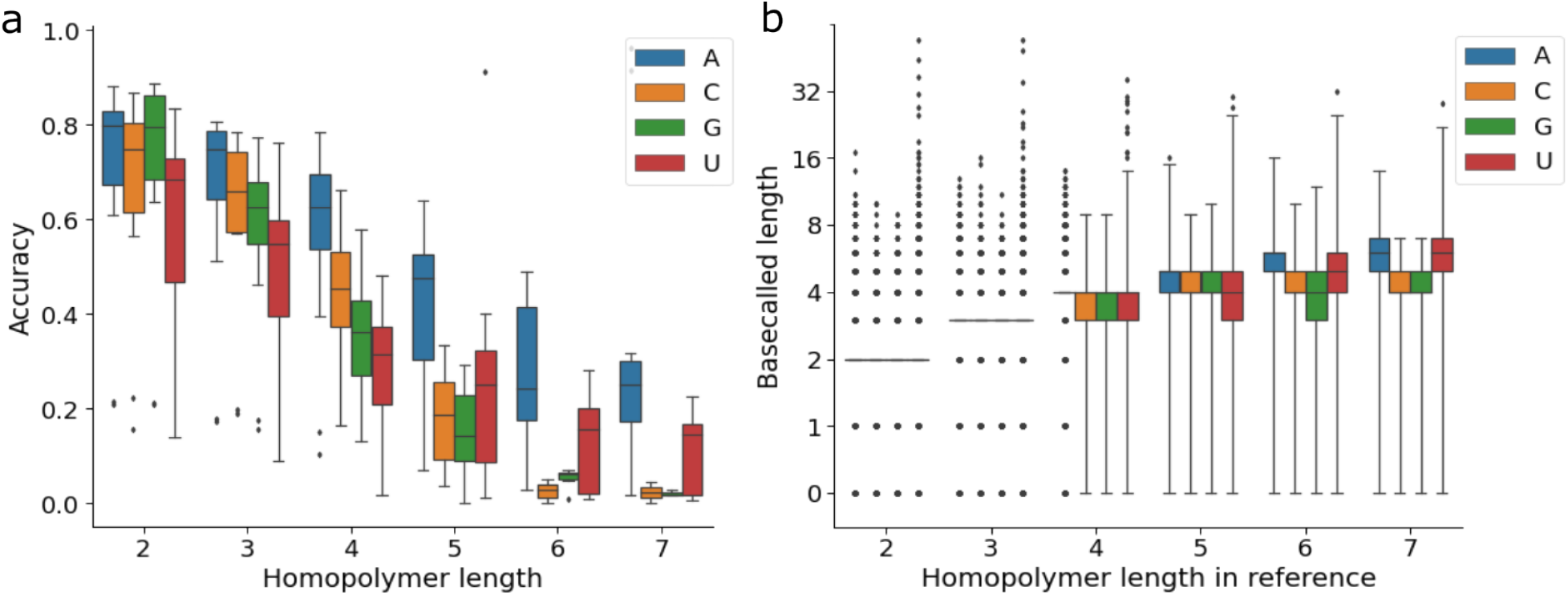
Systematic errors in homopolymeric regions. a) Basecalling accuracy of homopolymers with reference length between 2 and 7, grouped by nucleotides. b) Basecalled lengths of homopolymers with reference length between 2 and 7, when the basecalled motif is a homopolymer of the correct nucleotide type.

Looking at the specific nucleotides, A and U homopolymers are more accurate than Cs and Gs. Among the distinct bases, homopolymers of A are the most robustly basecalled with increasing lengths, being the only nucleotide with over 50% accuracy at length of 5. Interestingly, in nanopore DNA sequencing datasets [Delahaye and Nicolas, 2021], higher accuracies of A and T homopolymer motifs have also been observed. This is likely due to the larger presence of A/U homopolymers in the training data for basecallers arising from their overrepresentation in genomes and transcriptomes [Dechering et al., 1998].

### 3.4 Context-dependent sequencing errors

At each time step of nanopore sequencing, around five consecutive RNA bases reside in the pore simultaneously and produce a signal event, which, if distinct enough from the surrounding signals, will lead to a correct basecall. The surrounding sequence neighborhoods of a particular position, or the “sequence contexts”, are known to have a effect on the accuracy of the center. To examine the impact of sequence contexts on the accuracy in dRNA-seq data, we extended the analysis of single nucleotide errors by including the 3-mer sequence contexts – the errors of the center nucleotide, conditioned on that the neighboring 2 bases are correctly basecalled. We found strong biases in the error profiles of each nucleotide towards specific sequence contexts with a strong correlation between organisms (Figure 4a). Among the four RNA nucleotides, guanine is the most robust center base over its sequence contexts, with overall lower deletion than mismatch rate, indicating that the signal of G is more distinct than other nucleotides. The accuracy of cytosine is the most impacted by the sequence contexts: the presence of uridines in its neighborhood can drastically change its errors, with “UCU” having both the highest deletion and mismatch rates. Except for guanines, we did not observe a general correlation between mismatch and deletion errors, reaffirming that the two errors likely arise from different signal-level causes.

**Figure 4:**
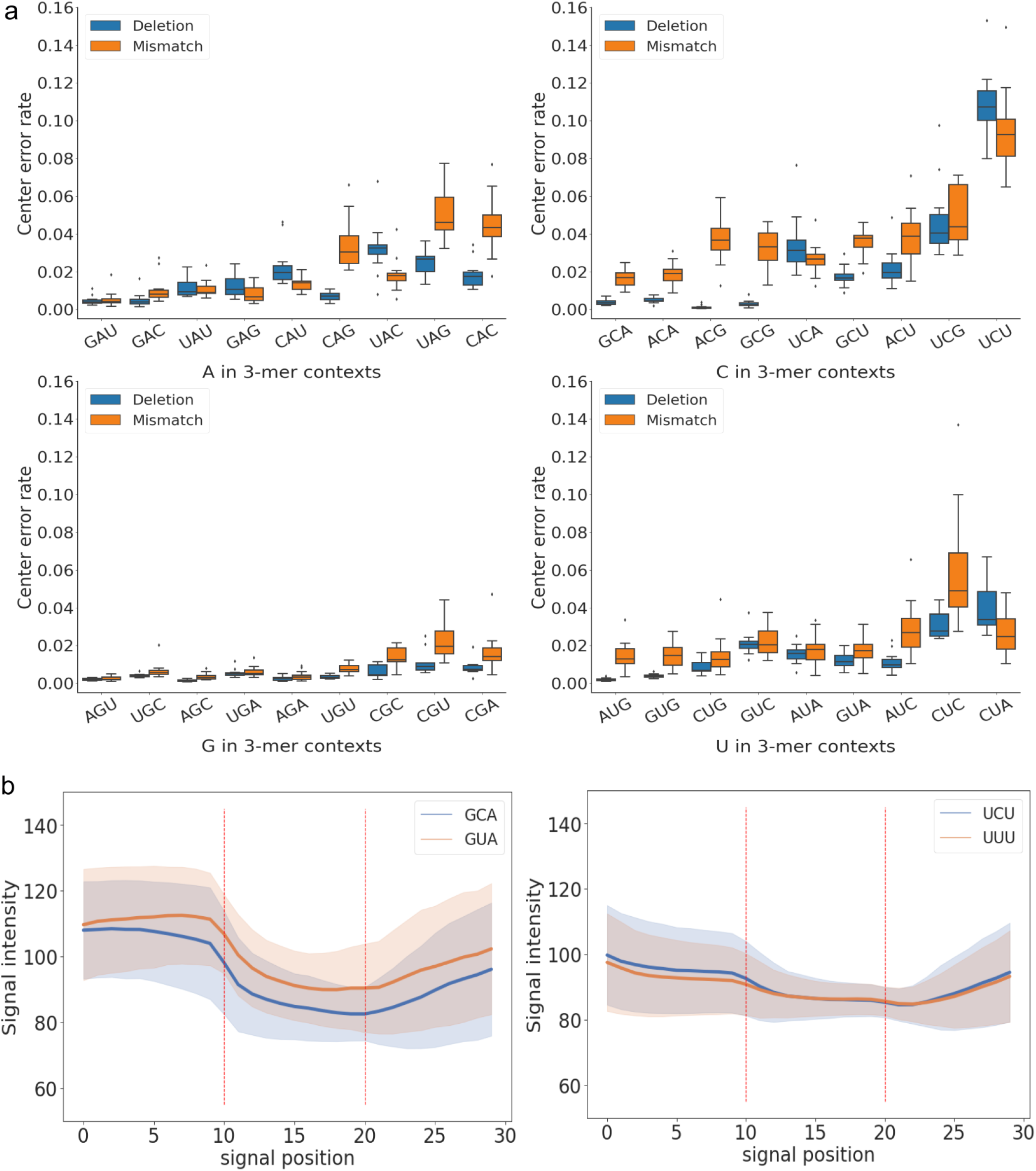
Context-dependent errors. a) The error profiles of the center nucleotides across 3-mer contexts, conditional on that the neighboring two bases are correctly basecalled. Each data point represents the mean error rate of all such 3-mer motifs in one dataset. b) The signal distribution of two pairs of motifs (GCA vs. GUA and UCU vs. UUU) at correctly basecalled positions. The concrete lines are the mean signal intensity and the shaded error bars are the mean signal standard deviation at each position. As the dwell times can differ, a running window is applied to extract the mean signal intensity with a step size of length of signals divided by 10, resulting in ten signal samples for each base. The red dotted lines represent the nucleotide boundaries at positions 10 and 20.

To exemplify the signal-level sources of context-dependent errors, we showcase the signal plots of two 3-mer motifs, GCA and UCU, which are respectively the most and least accurate 3-mer centered at cytosine, and their closest mismatched counterparts, GUA and UUU (Figure 4b). The differences at the center position of UCU and UUU in signal intensity and standard deviation are minimal, whereas GCA and GUA motifs show more distinct separation in signal space at all three positions. A likely cause could be that in the absence of distinct signals (in the case of UCU and UUU), the basecaller would have to rely on longer range information to determine the nucleotide identity at the center, resulting in increased complexity and errors.

### 3.5 Estimating read errors from read and base quality scores

The basecaller-generated quality scores are a helpful estimate for read accuracy without relying on the availability of reference genomes. There are two types of quality scores provided by ONT basecallers such as Guppy, 1) read-level quality scores (Q-scores) in the `sequence_summary.txt` file, and 2) base-level quality scores encoded as ASCII characters found in *fastq* files for each base on each read. Both types of quality scores are Phred quality scores, e.g. Q10 is equivalent to an error rate of 10% and Q20 equivalent to 1% (Equation 1). For dRNA-seq basecalling with Guppy, the default read Q-score filter is set to 7 (corresponding to an expected read accuracy of 80%). Reads with a lower Q-score than 7 are stored in the *failed* folder and are typically not included in downstream analysis.

To evaluate the accuracy of basecaller-estimated error rates, we compared the estimated error rate from read Q-scores with the empirical read error rates measured after alignment. For most organisms, the measured read error rates can be reasonably estimated at read Q-scores higher than 10, while at lower Q-scores the read error is often overestimated (Figure 5a). This suggests that the default Q-score filter of 7 may be overly stringent. One important note here is that the measured read errors are a result of both basecalling and alignment. That is, aligners such as minimap2 employ “soft clipping” and do not always align reads at full length. Therefore, the empirical read error rates only reflect the errors at the aligned part of the reads. Indeed, reads with lower Q-scores tend to have more soft-clipped bases (Figure 5b), indicating a larger number of errors at the start or end of the reads.

**Figure 5:**
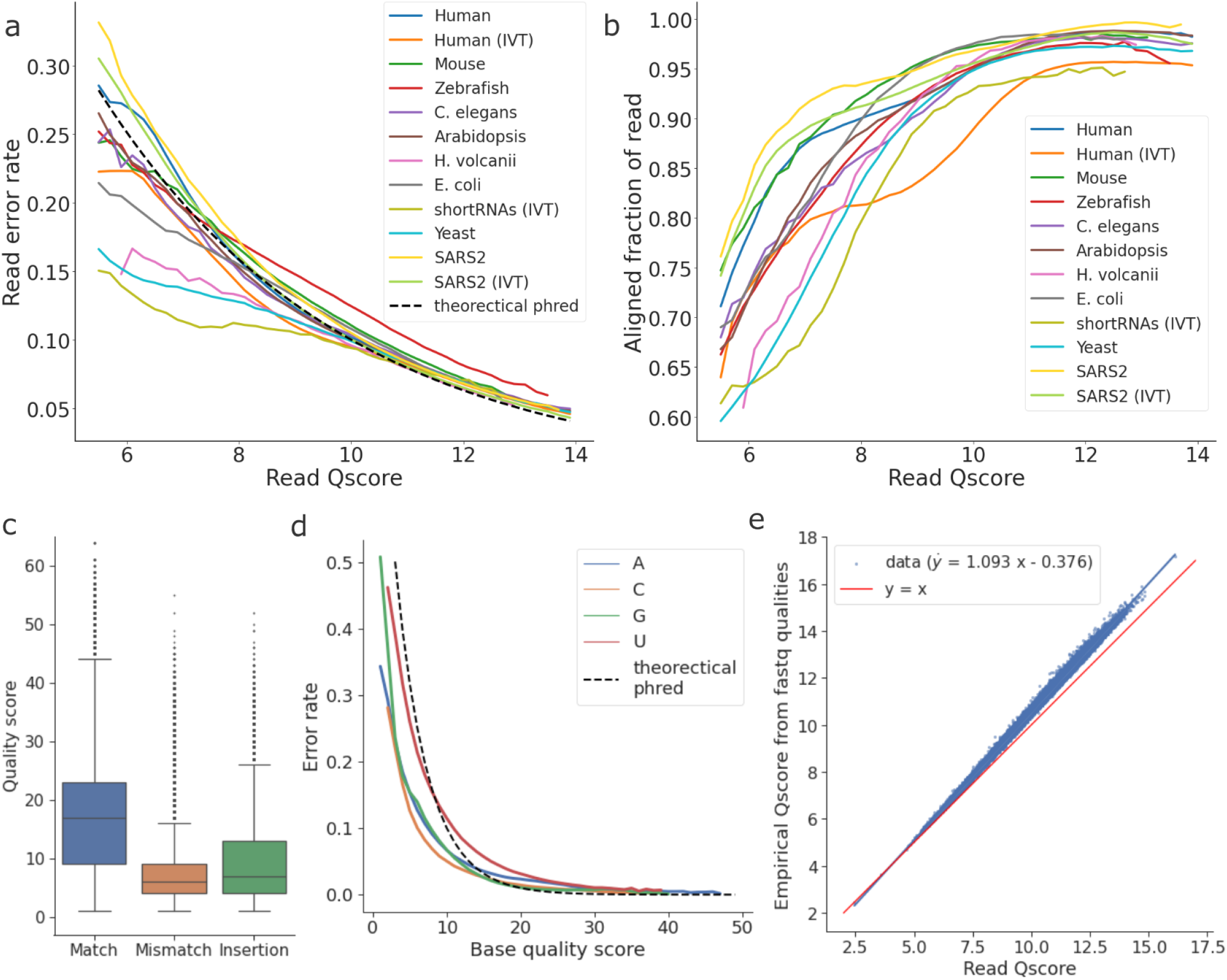
Read and base quality scores in dRNA-seq. (a) The relationship between read Q-scores and read error rates. The dashed line reflects the theoretical error rate based on the Phred quality scores. (b) The relationship between read Q-scores and the aligned fraction of reads after alignment for each dataset. (c) The distribution of per base quality scores, grouped by the per-base error type. (d) The relationship between per base quality scores and per-base error rate (mismatch rate + insertion rate), grouped by nucleotide. (e) The relationship between read Q-scores and “empirical” Q-scores computed from per base quality scores of reads in *fastq* files, based on the native human sample. The linear model is fitted by the ordinary least square method.

In addition to the read Q-scores, the base-level quality scores for each basecalled read can be useful in predicting per base errors in the basecalls (mismatches and insertions). We found that correctly sequenced bases tend to have higher quality scores than mismatched and inserted positions (Figure 5c). Moreover, except for uracil, the per-base error rate can be accurately estimated when base-level quality scores are higher than 15 (Figure 5d).

Theoretically, the read Q-scores are obtained by the log transformation of the mean per base error estimate, based on the relationship described in Equation 2. However, we observed a systematic deviation between the empirically computed Q-scores and the original read Q-scores reported by Guppy, especially at higher values (Figure 5e). This deviation is found across all datasets and could be a consequence of the loss of numerical precision in the discretisation of per-base quality scores into ASCII characters. Nevertheless, the deviation remains highly linear (fitted with an ordinary least square model, p-value *≈* 0).

### 3.6 Adapter detection failure

Whilst read lengths are not found to impact read accuracy (Supplementary Figure S1), we observed that the discrepancy between the basecalled read length and the aligned read length becomes larger as read length decreases (Figure 6a), as a consequence of more soft-clipped bases at the start and end of the reads.

**Figure 6:**
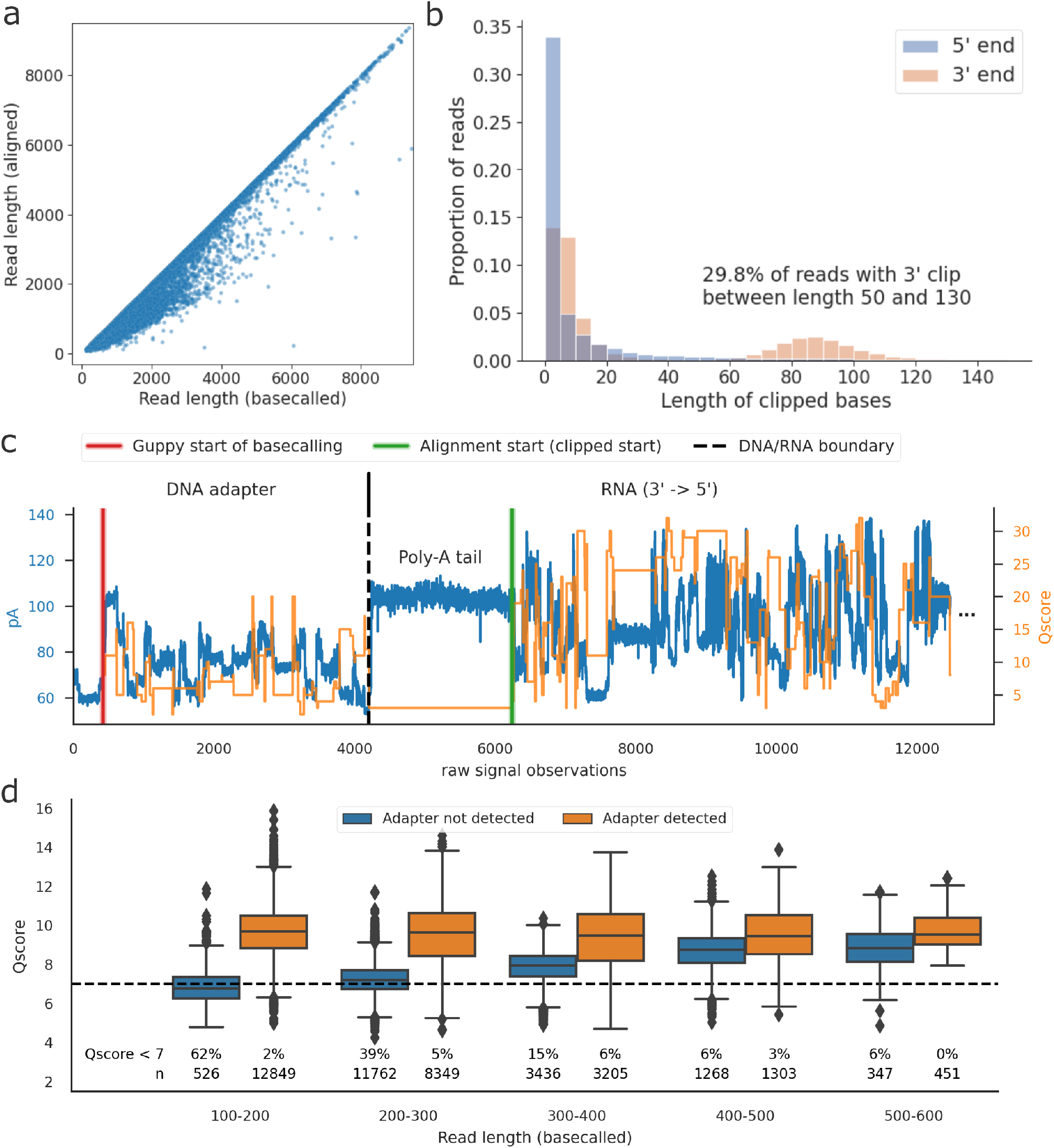
Adapter detection failure in Guppy. a) The relationship between the basecalled read length and the aligned read length after soft-clipping, based on the native human sample. b) The length distribution of the soft-clipped bases at the 5’ and 3’ ends of reads, the latter of which shows a clear bimodal distribution, indicative of adapter detection failure, based on the native human sample. c) A signal visualization of a read whose adapter failed to be detected by Guppy. The left y-axis shows the signal intensity value (in pA) and the right y-axis shows the quality scores per base. d) The distribution of read Q-scores depending on whether the adapter was detected successfully, grouped by read length, based on the short RNA dataset. Adapter detection failure can reduce Q-score for short reads, leading to them being filtered at the conventional Q-score threshold of 7 (dashed line).

We then counted the soft-clipped bases per read and found that in contrast to the 5’ end, the length distribution of the soft-clipped bases at the 3’ end showed a clear bimodal pattern, with a second mode locat-ing roughly between 50 and 130 bases and accounting for 30% of the reads (Figure 6b, native human). The unique bimodality of 3’ soft-clips are found across datasets, with the second mode accounting for up to 41.5% of the reads in the zebrafish sample (Supplementary Figure S4), indicating that this is a common pattern in dRNA-seq data.

Upon inspecting the signal data of these reads, we dis-covered that the soft–clipped bases at the 3’ end were actually erroneous basecalls originating from the DNA adapter. For these reads, Guppy began basecalling at the beginning of the adapter, instead of at the start of the RNA segment. The beginning of reads with this issue typically contains a jump in signal intensity that is comparable to the one typically seen at the end of the adapter, marking the DNA/RNA boundary (Figure 6c).

To evaluate the impact of adapter detection failure on the quality of downstream RNA basecalls, we compared the alignment error rates and basecall quality scores of reads with and without successful adapter detection. We found that basecalls in the DNA adapter are of low quality (Supplementary Figure S5a), but we did not observe that having these spurious basecalls at the start of the read negatively impacts the error rate of the subsequent RNA basecalls (Supplementary Figure S5b). However, for shorter reads (less than 400 bases), we observed that the spurious DNA basecalls significantly impact the read Q-score, dragging it below the default Q-score cutoff of 7 and leading to unnecessary data loss (Figure 6d, Supplementary Figure S5c).

### 3.7 Comparison with the community basecaller RODAN

For dRNA-seq data, RODAN [Neumann et al., 2022] is currently the only non-ONT basecaller published by the research community. To examine whether RODAN has significantly improved over Guppy in base-calling performance, we re-basecalled all the datasets in this study with RODAN and compared its performance with Guppy (Supplementary Table S2). Among the datasets, four organisms (native human, *C. elegans, E. coli*, and *Arabidopsis*) were included in the training data of RODAN. RODAN performed 2–5% better than Guppy in three of these four datasets (except *Arabidopsis*). In the species outside its training set, RODAN has a 1–2% higher median read accuracy than Guppy on the mouse, *H. volcanii*, the short RNAs and the IVT SARS2 datasets; however, on the zebrafish dataset it has a 4.8% lower median read accuracy.

While the read accuracies are comparable on most datasets, the reads from RODAN have shorter median aligned read lengths in 10 out of 12 datasets (Figure 7a, Supplementary Table S2). The largest drop in the aligned read lengths is observed in the native SARS2 dataset, where the median read length from RODAN is 708 nucleotides shorter, i.e. a 34% decrease. To explore whether RODAN’s performance is related to read length, we grouped reads by length and found that RODAN performs overall better than Guppy in shorter reads, but the improvement diminishes as read length increases, with the exception of organisms in RODAN’s training dataset (Figure 7b).

**Figure 7:**
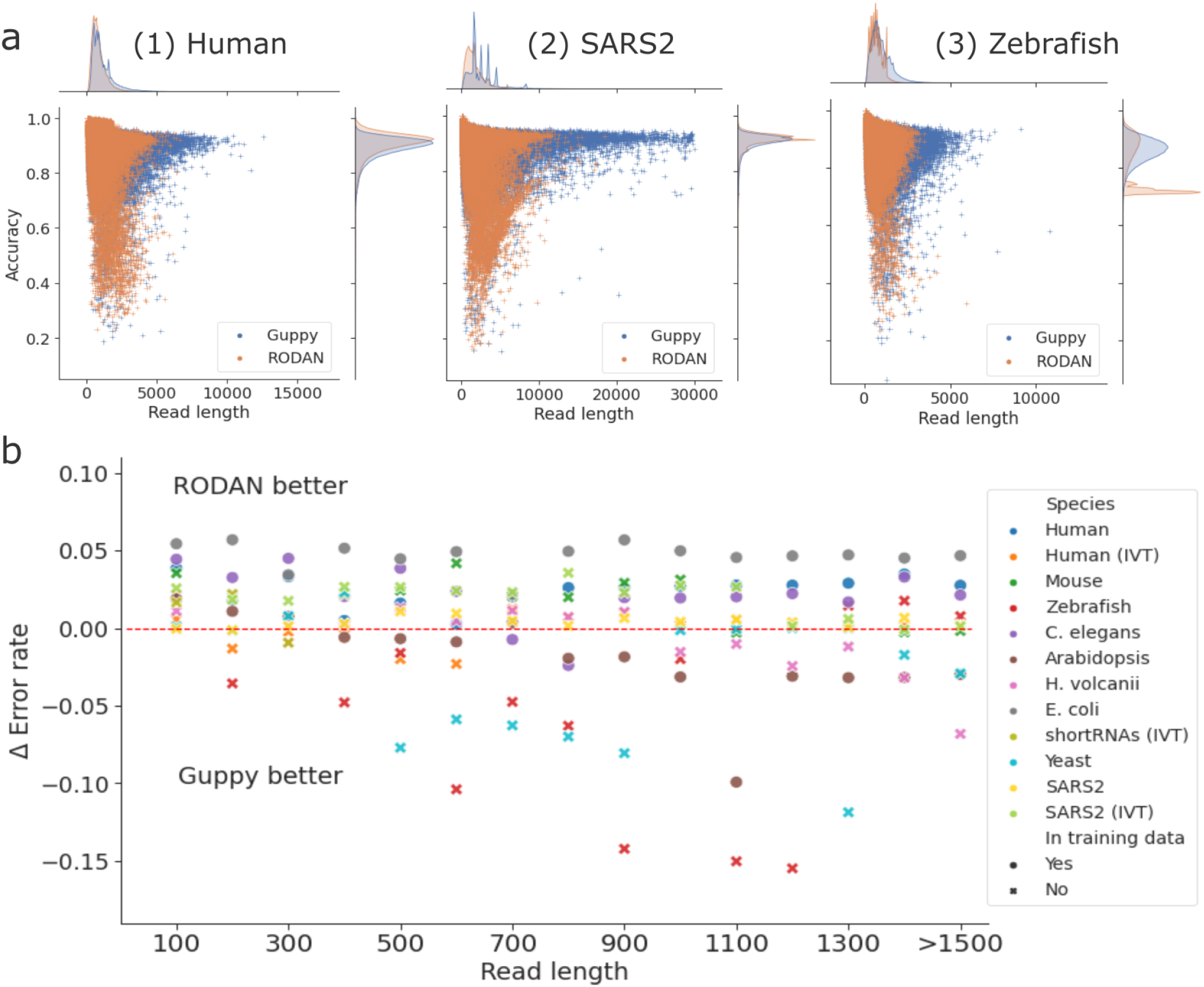
Comparison of basecalling performance between Guppy and the community basecaller RODAN. (a) The joint and marginal distributions of read length and read accuracy in 1) the native human, 2) SARS2 and 3) zebrafish samples. Each dataset was subsampled to 200,000 reads due to the runtime for kernel density estimation. (b) The relationship between aligned read length and the performance difference of RODAN versus Guppy. The Δ Error rate is Error_Guppy_ - Error_RODAN_, thus a positive value means RODAN is more accurate than Guppy. The symbols represent whether the dataset was present (•) or absent (×) in the training set of RODAN.

Lastly, the systematic sequencing errors at single nucleotide and motif levels are also prevalent in the reads basecalled by RODAN (Supplementary Figure S7), suggesting that the fundamental causes of sequencing errors reported in this study are not basecaller-specific, but could rather be related to intrinsic difficulties imposed by the raw signal data.

## 4 Discussion

Since the release of the first MinION device by ONT, the accuracy of nanopore DNA sequencing has significantly improved due to developments in both pore chemistry and basecalling algorithms [Rang et al., 2018, Sereika et al., 2022]. At the same time, dRNA-seq has gained considerable popularity due to its reduced bias and the ability to sequence RNA base modifications and characterise the epitranscriptome. However, improvements in the sequencing accuracy of dRNA-seq has so far been limited [Jain et al., 2022]. Here, we systematically evaluated the accuracy and systematic error patterns of dRNA-seq in datasets covering diverse organisms across the tree of life. We found that the read accuracy is around 90% across organisms in both native and IVT samples, and that deletions account for the majority of errors.

The high error rate of dRNA-seq has several important implications in downstream analyses. Firstly, as each native RNA read represents a single polyadenylated RNA transcript that is potentially protein coding, one promising application of dRNA-seq is to identify and quantify the expression of novel transcript isoforms and open reading frames (ORFs). However, the high insertion and deletion (indel) error rates can often lead to frameshift errors in gene prediction, which is a well-known challenge in long-read DNA-seq [Watson and Warr, 2019]. To address this, Depledge et al. [2019] attempted to reduce errors by correcting dRNA-seq reads with Illumina short reads, yet the remaining indel errors still precluded ORF prediction in more than 80% of the reads. Moreover, despite being much longer than Illumina reads, alignments of dRNA-seq reads to transcriptomes often produce a considerable amount of secondary alignments with similar mapping scores to primary alignments, thus making the identification of true transcript origin difficult [Soneson et al., 2019, Price et al., 2020], which we suspect to also be a result of the high error rate.

In addition to the overall error rates, we also observed systematic errors across our datasets that show sequence-specific patterns and biases. The systematic errors at single nucleotide and motif levels are particularly relevant to the application of dRNA-seq in characterising the epitranscriptome by detecting native RNA base modifications. Current computational methods for detecting RNA modifications attempt to identify potentially modified sites through either higher basecalling error rates or deviations in the signal at a particular position, by comparing with control samples that have fewer modifications (in the case of writer gene knockouts) or no modifications (in the case of IVT samples) [Begik et al., 2022b]. Because there are inherent background errors and signal changes even in unmodified RNA, the inclusion of control samples is necessary to reduce the false positive rate of modification detection [Kim et al., 2020], thus limiting our ability to detect rare modifications *de novo* when negative controls are hard to obtain.

It is worth noting that we did not investigate the role of native base modifications on the observed error patterns. While the modification-free IVT human dataset has less errors than its native counterpart, the same pattern is not seen in the two SARS2 datasets. Certain types of modifications, such as m^6^A, m^5^C and pseudouridine, have been found to lead to an overall increase in basecalling errors, but the error patterns are also dependent on the local sequence contexts [Parker et al., 2020, Leger et al., 2021]. Moreover, one would expect that commonly modified motifs, such as the DRACH motif of m^6^A, are already present in the training data and can be learned by the basecaller, but less so for the rare or species-specific types of modifications. Indeed, a careful investigation of the differences between the native and the unmodified IVT samples is warranted to uncover the likely complex relationship between basecalling errors and specific types of modifications.

In general, there are two sources from which sequencing errors in nanopore sequencing data can arise: firstly, the raw signals produced by the nanopore sequencer and secondly, the basecalling algorithm translating the signal data into nucleotide sequences [Rang et al., 2018]. In addition to improvements of ONT devices and chemistry for nanopore DNA-seq, there is also continuous development of new basecallers released by both ONT and the broader research community [Wick et al., 2019, Silvestre-Ryan and Holmes, 2021]. To facilitate such efforts, ONT provided soft-wares such as the *Taiyaki* tool (https://github.com/nanoporetech/taiyaki) to support the development of research basecallers. However, for dRNA-seq data, RODAN [Neumann et al., 2022] remains currently the only published community basecaller outside of ONT. From our evaluation, the performance of RODAN holds up well against Guppy in terms of read accuracy, especially for organisms in its training data. As species-specific training data are known to improve performance of nanopore basecallers in those species [Wick et al., 2019, Ferguson et al., 2022], the improvements of RODAN suggest a promising direction for training species-specific basecallers also for dRNA-seq data. Lastly, the presence of the same systematic error patterns in RODAN points to more fundamental causes of errors in the raw signal data, necessitating further development of pore chemistry to improve the decoding of raw signal data.

In conclusion, while dRNA-seq offers exciting opportunities for studying RNA transcription and modifications, our evaluation highlights the need for continued improvements in data quality and accuracy. Clearly, further development and optimization of dRNA-seq protocols, pore chemistry and basecalling algorithms are desirable. At the same time, appropriate computational methods for data quality control, uncertainty quantification, and error correction are needed to mitigate the effects of high error rates and systematic biases in downstream analyses, especially in the area of *de novo* transcript identification and RNA-modification detection.

## Data availability

The code and scripts to reproduce the analyses and plots are available on https://github.com/KleistLab/nanopore_dRNAseq.

## 5 Supplementary materials

**Table S1:** Details of the datasets. Statistics regarding the aligned reads (counts, quality scores and lengths) and their accuracies of each dataset included in this study.

| Dataset | Reads aligned | med. Q-score | med. Length | Acc | Mis | Ins | Del |
| --- | --- | --- | --- | --- | --- | --- | --- |
| Human | 1483755 | 10.2 | 876 | 0.901 | 0.027 | 0.026 | 0.045 |
| Human (IVT) | 2015950 | 10.7 | 486 | 0.916 | 0.022 | 0.022 | 0.039 |
| Mouse | 181977 | 9.6 | 616 | 0.878 | 0.036 | 0.025 | 0.059 |
| Zebrafish | 553604 | 9.6 | 839 | 0.867 | 0.037 | 0.020 | 0.073 |
| C. elegans | 227372 | 11.0 | 688 | 0.915 | 0.020 | 0.019 | 0.044 |
| Arabidopsis | 1010943 | 10.7 | 878 | 0.911 | 0.022 | 0.021 | 0.045 |
| H. volcanii | 22315 | 9.9 | 505 | 0.905 | 0.028 | 0.022 | 0.043 |
| E. coli | 198448 | 9.3 | 662 | 0.876 | 0.040 | 0.032 | 0.049 |
| shortRNAs (IVT) | 45028 | 8.5 | 160 | 0.899 | 0.026 | 0.019 | 0.051 |
| Yeast | 324889 | 9.4 | 314 | 0.898 | 0.024 | 0.024 | 0.050 |
| SARS2 | 575673 | 11.0 | 2078 | 0.914 | 0.022 | 0.020 | 0.044 |
| SARS2 (IVT) | 2316635 | 10.5 | 1572 | 0.908 | 0.024 | 0.023 | 0.044 |

**Figure S1:**
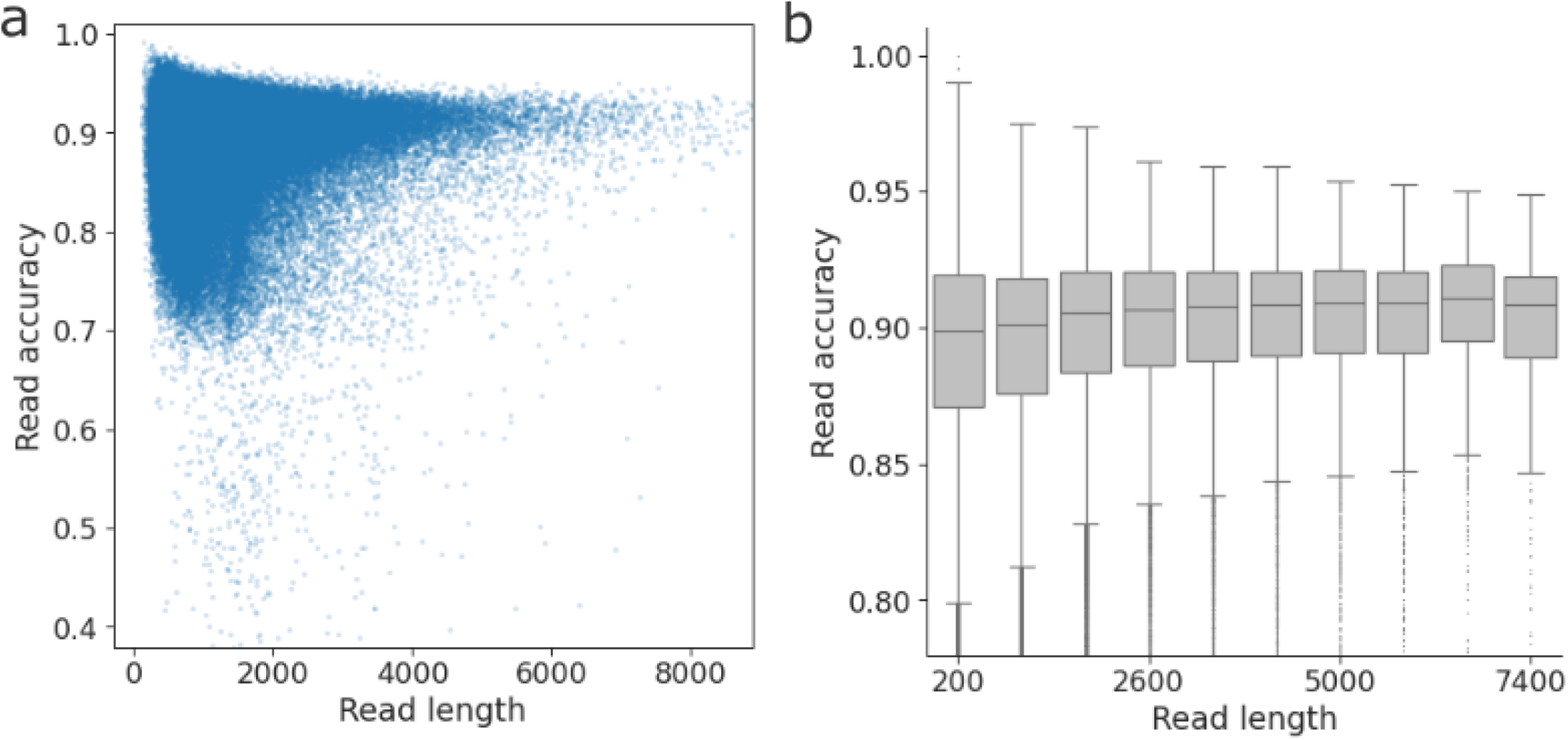
The relationship between read length and read accuracy. a) Scatter plot of read length versus read accuracy, based on the native human dataset. b) A boxplot of the same data, grouped by read length.

**Figure S2:**
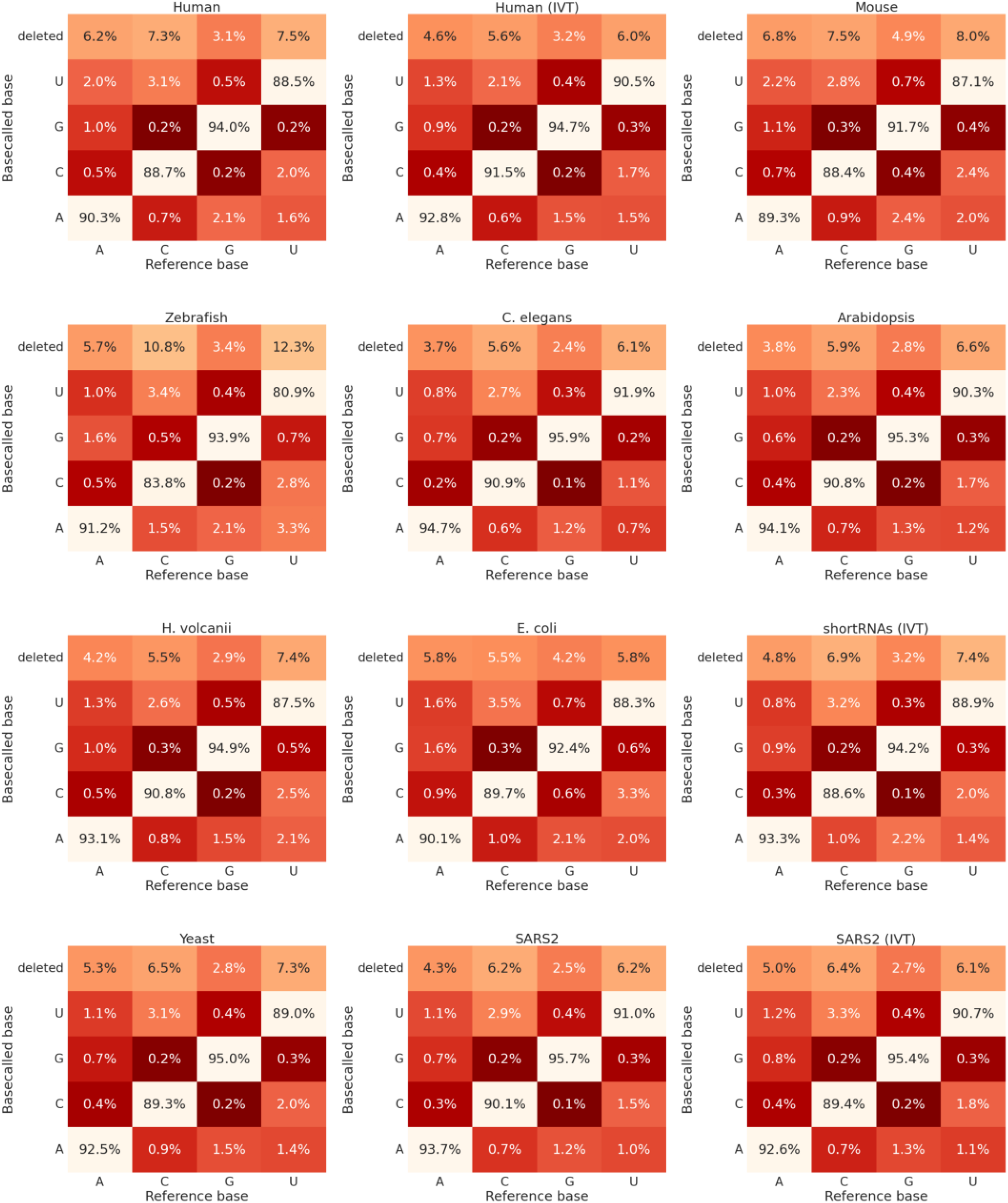
Single nucleotide error profiles across organisms. Confusion matrix showing the proportion of each base in reference being correctly basecalled, miscalled or deleted, for each organism included in the study.

**Figure S3:**
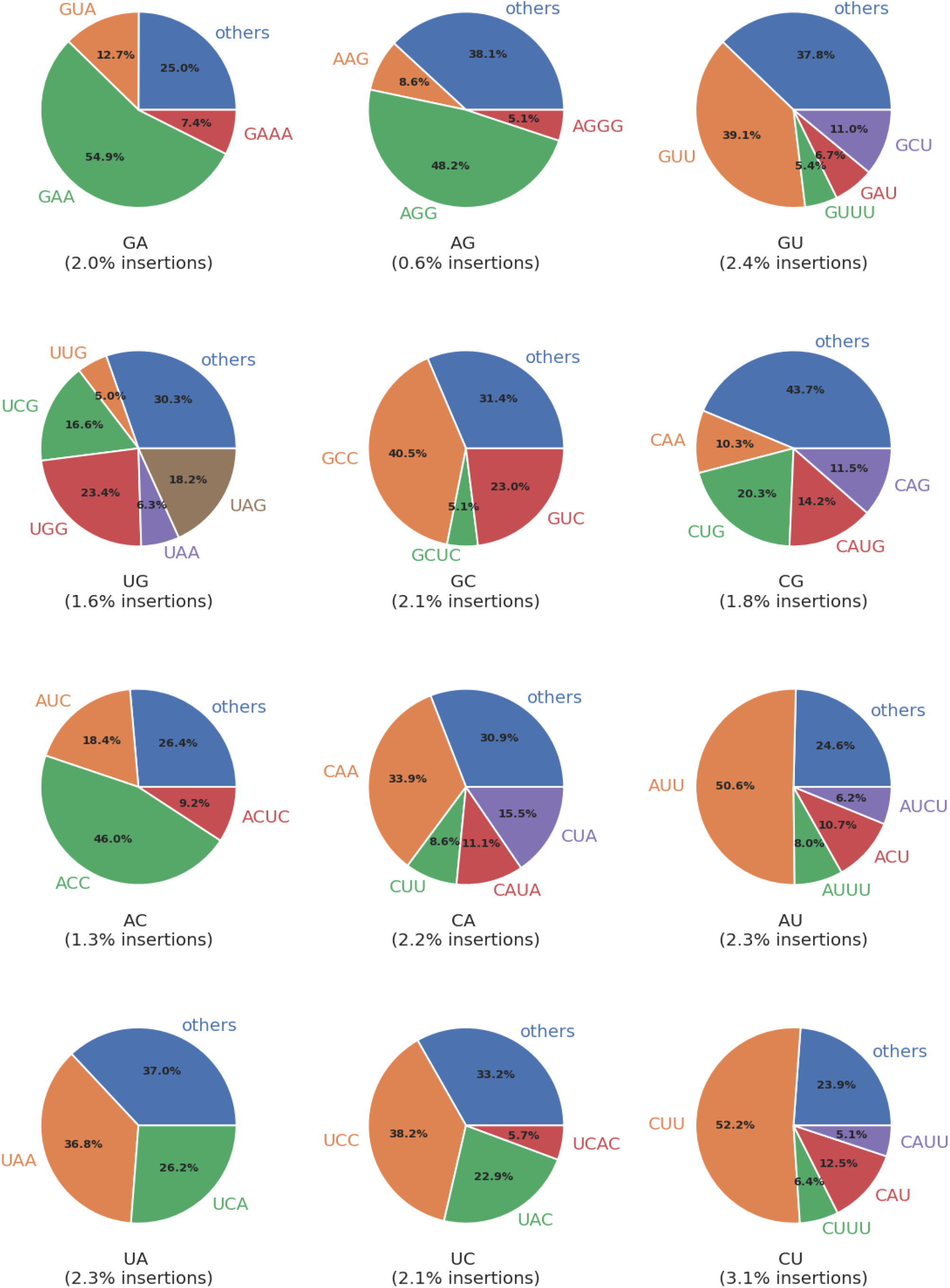
Insertion profiles of 2-mer motifs. Pie charts showing the proportion of different insertion errors, based on the native human sample. Inserted motifs fewer than 5% of the total number of insertions are grouped into “others”.

**Figure S4:**
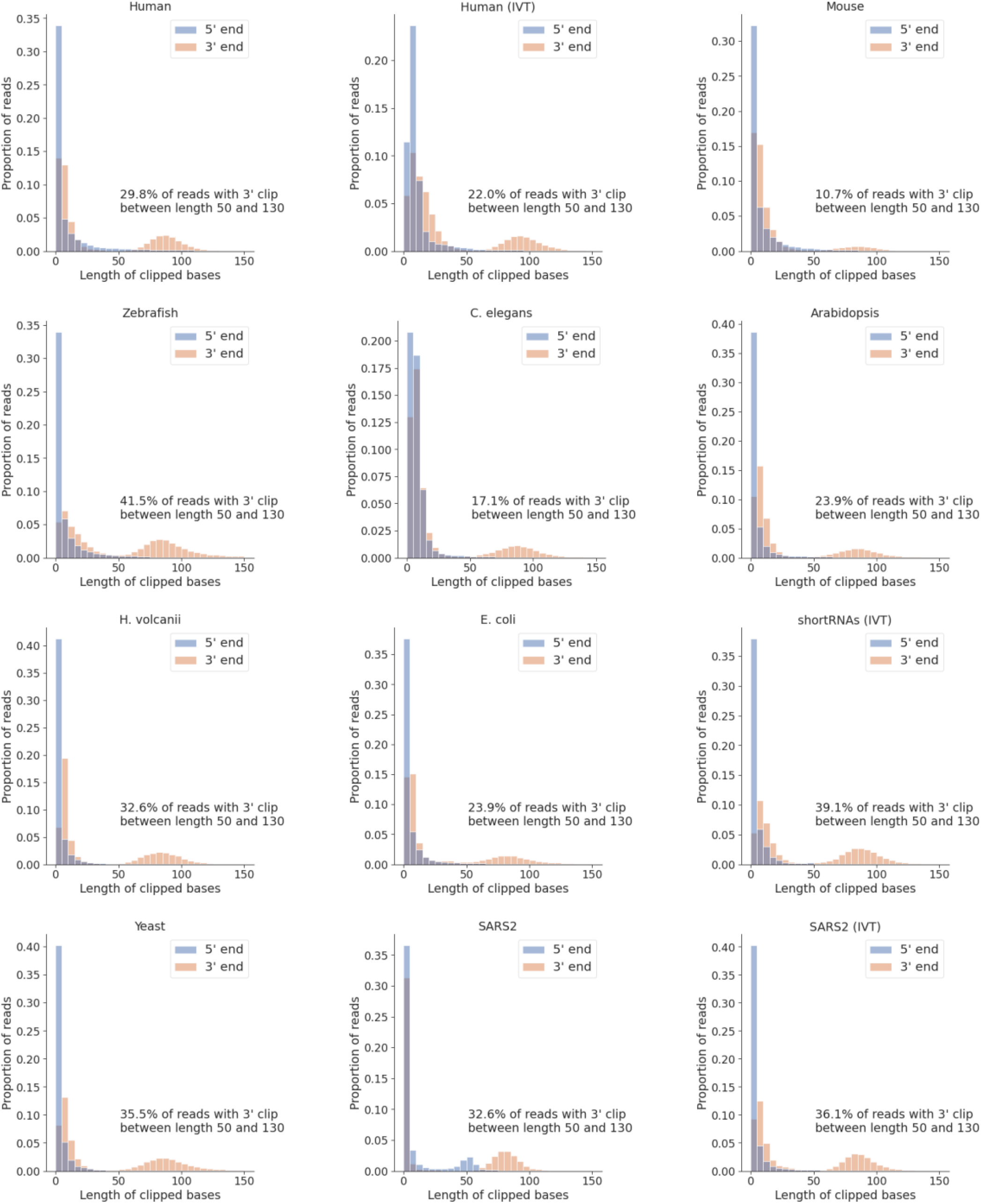
Bias in soft-clipped bases at the 3’ end. The length distribution of the soft-clipped bases at the 5’ and 3’ ends of reads in each dataset in this study.

**Figure S5:**
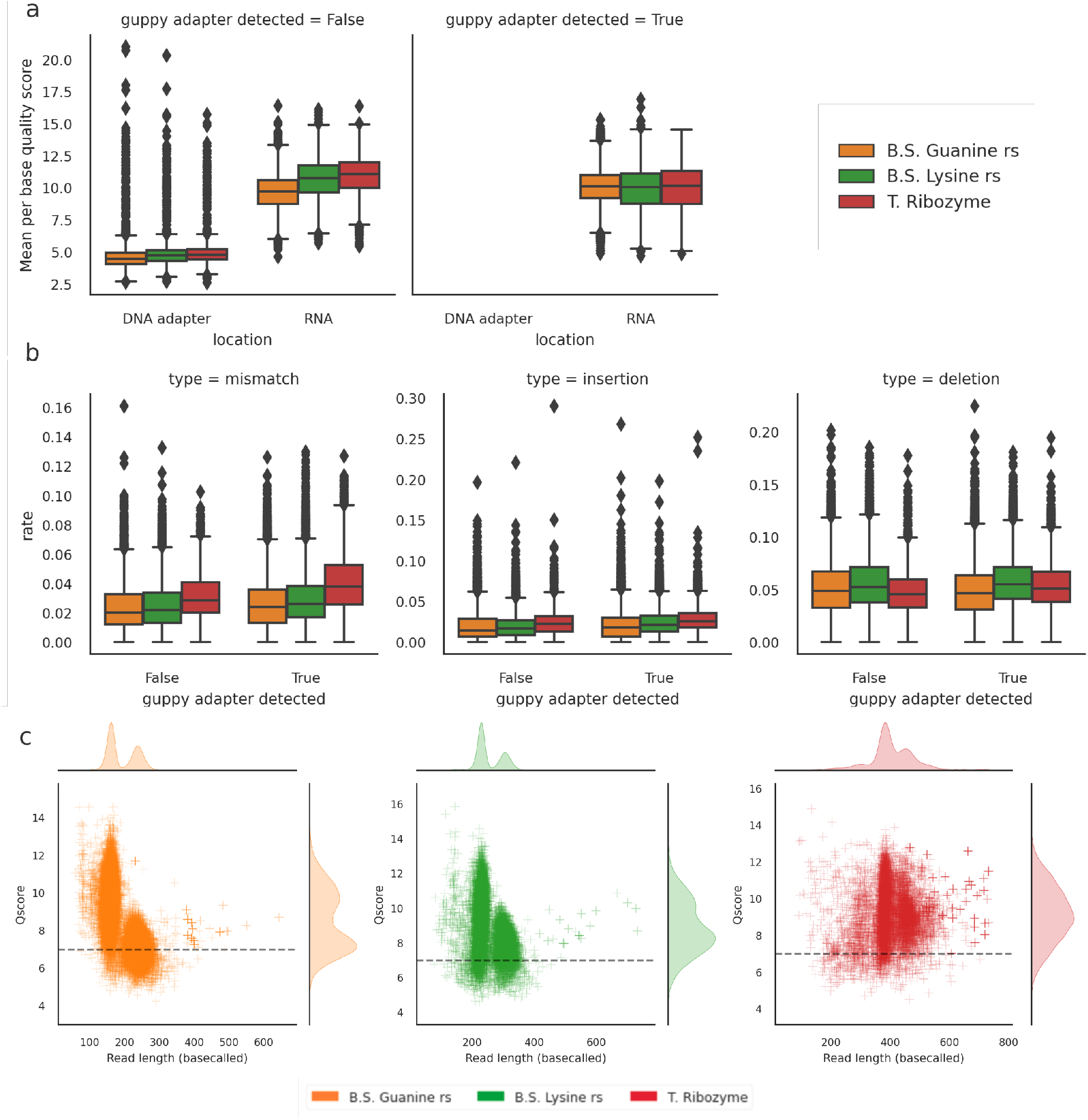
Comparison of reads with and without successful adapter detection. a) Basecalls in the DNA adapter are of low quality, but adapter detection failure does not appear to affect the quality scores of the subsequent RNA basecalls. b) Failure to detect the adapter does not affect the error rates of the RNA basecalls after alignment. c) The impact of adapter detection failure on basecalled read length and read Q-score in short reads. Adapter detection failure causes bimodality in basecalled read length. The spurious DNA basecalls can significantly impact the read Q-score, potentially leading to data loss. The dashed line represents the conventional filter value of 7. Plots are based on the IVT short RNA dataset.

**Table S2:** Basecalling performance of RODAN versus Guppy across organisms. The differences in each metric from Guppy are indicated in brackets.

| Dataset | In training data | Reads aligned | med. Length | Acc | Mis | Ins | Del |
| --- | --- | --- | --- | --- | --- | --- | --- |
| Human | yes | 1648051<br>(+164296) | 725<br>(-151) | 0.923<br>(+0.022) | 0.019<br>(-0.008) | 0.021<br>(-0.004) | 0.035<br>(-0.009) |
| Human (IVT) | no | 1729393<br>(-286557) | 375<br>(-111) | 0.913<br>(-0.003) | 0.023<br>(+0.001) | 0.027<br>(+0.005) | 0.037<br>(-0.002) |
| Mouse | no | 85286<br>(-96691) | 591<br>(-25) | 0.903<br>(+0.025) | 0.018<br>(-0.017) | 0.015<br>(-0.01) | 0.059<br>(-0.0) |
| Zebrafish | no | 155285<br>(-398319) | 659<br>(-180) | 0.818<br>(-0.048) | 0.051<br>(+0.014) | 0.018<br>(-0.002) | 0.114<br>(+0.041) |
| C. elegans | yes | 222931<br>(-4441) | 558<br>(-130) | 0.941<br>(+0.026) | 0.012<br>(-0.008) | 0.014<br>(-0.005) | 0.031<br>(-0.013) |
| Arabidopsis | yes | 1097675<br>(+86732) | 471<br>(-407) | 0.903<br>(-0.008) | 0.032<br>(+0.011) | 0.028<br>(+0.007) | 0.033<br>(-0.012) |
| H. volcanii | no | 14258<br>(-8057) | 959<br>(+454) | 0.916<br>(+0.011) | 0.028<br>(-0.0) | 0.035<br>(+0.013) | 0.021<br>(-0.022) |
| E. coli | yes | 506596<br>(+308148) | 578<br>(-84) | 0.921<br>(+0.045) | 0.025<br>(-0.015) | 0.023<br>(-0.01) | 0.026<br>(-0.023) |
| shortRNAs (IVT) | no | 41329<br>(-3699) | 170<br>(+10) | 0.919<br>(+0.02) | 0.020<br>(-0.005) | 0.026<br>(+0.007) | 0.027<br>(-0.024) |
| Yeast | no | 372364<br>(+47475) | 297<br>(-17) | 0.892<br>(-0.006) | 0.028<br>(+0.004) | 0.026<br>(+0.002) | 0.060<br>(+0.01) |
| SARS2 | no | 858189<br>(+282516) | 1370<br>(-708) | 0.916<br>(+0.002) | 0.019<br>(-0.003) | 0.017<br>(-0.003) | 0.041<br>(-0.003) |
| SARS2 (IVT) | no | 2452303<br>(+135668) | 1103<br>(-469) | 0.920<br>(+0.012) | 0.018<br>(-0.005) | 0.021<br>(-0.002) | 0.033<br>(-0.011) |

**Figure S6:**
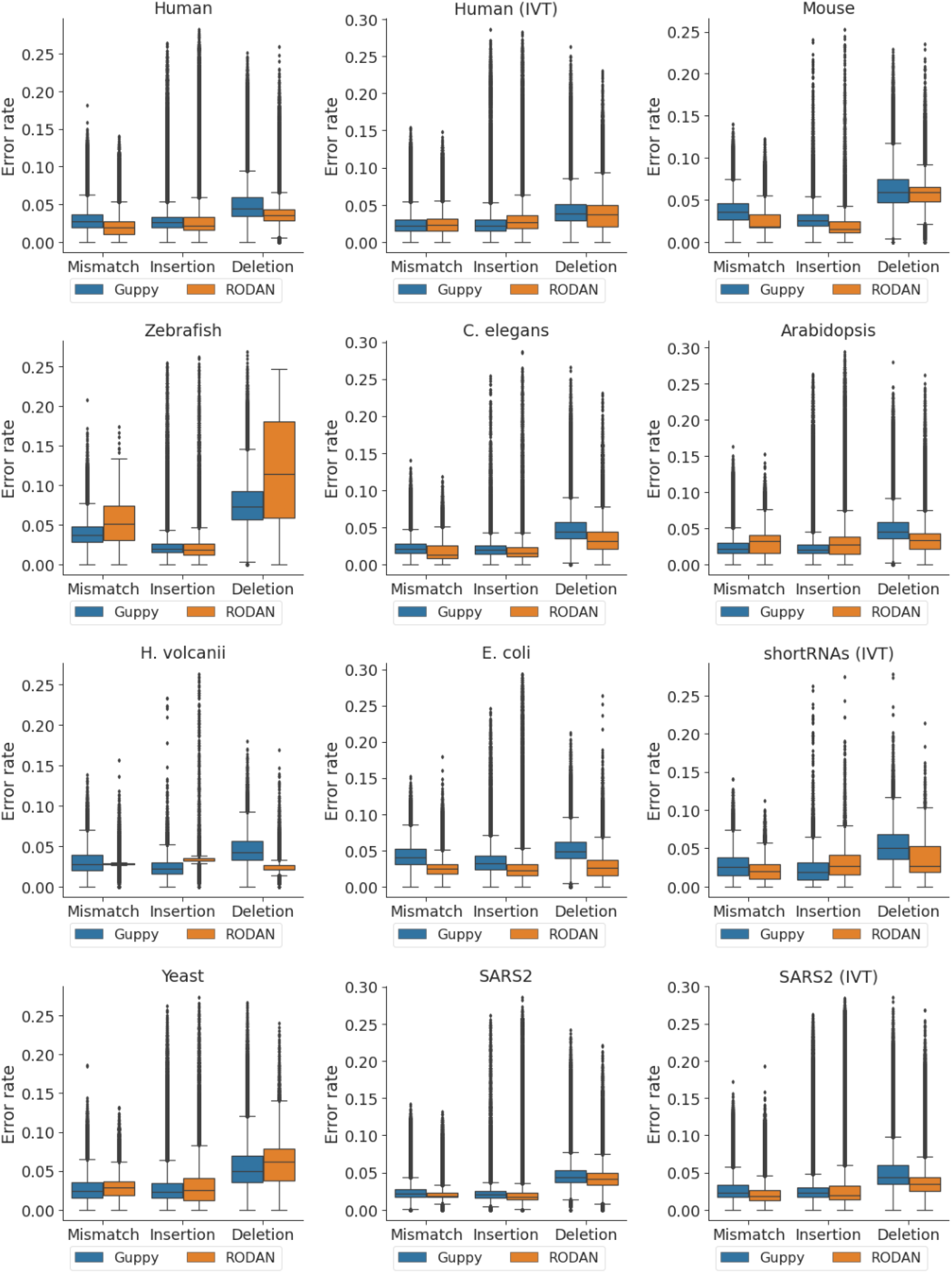
Comparison of basecalling errors between RODAN and Guppy. The mismatch, insertion and deletion rates for each dataset. Reads with accuracy lower than 70% (or equivalently, more than 30% errors), are filtered for visualization.

**Figure S7:**
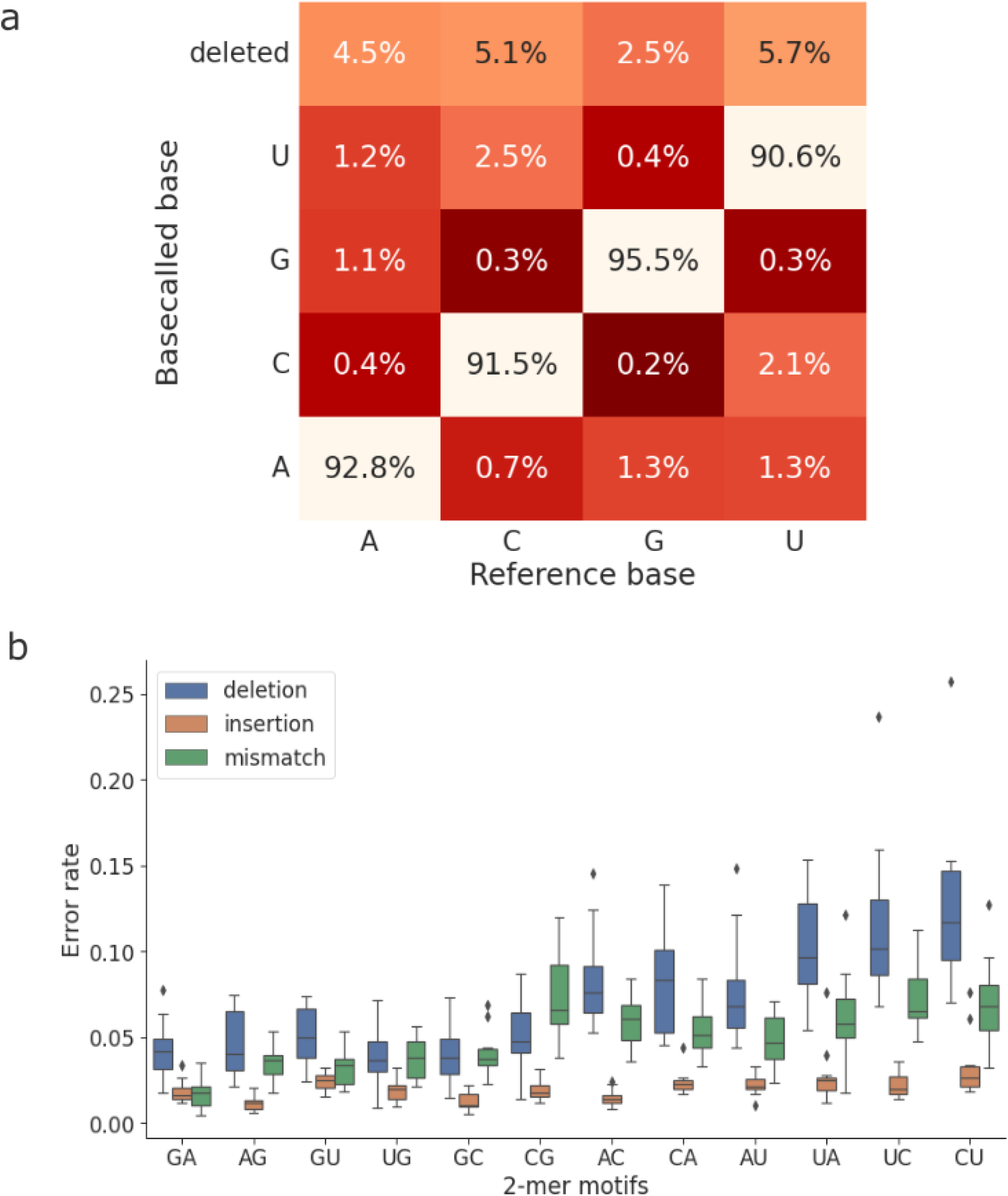
Systematic errors in RODAN. a) Confusion matrix showing the frequencies of each base being correctly basecalled, miscalled or deleted, computed by taking the mean of all samples basecalled by RODAN. b) The error rate of 2-mer motifs across the samples basecalled by RODAN, grouped by error type. Each data point corresponds to the mean error rate of the motif in one dataset.

